# Mouse models for pancreatic ductal adenocarcinoma are affected by the cre-driver used to promote KRAS^G12D^ activation

**DOI:** 10.1101/2023.09.12.557383

**Authors:** Fatemeh Mousavi, Joyce Thompson, Justine Lau, Nur Renollet, Mickenzie B. Martin, Jake McGue, Timothy Frankel, Parisa Shooshtari, Christopher L. Pin, Filip Bednar

## Abstract

The fundamental biology of pancreatic ductal adenocarcinoma has been greatly impacted by the characterization of genetically modified mouse models that allow temporal and spatial activation of oncogenic KRAS (KRAS^G12D^). The most commonly used model involves targeted insertion of a *cre* recombinase into the *Ptf1a* gene. However, this approach disrupts the *Ptf1a* gene, resulting in haploinsufficiency that likely affects sensitivity to oncogenic KRAS (*KRAS^G12D^*). The goal of this study was to determine if *Ptf1a* haploinsufficiency affected the acinar cell response to *KRAS^G12D^* before and after induction of pancreatic injury. We performed morphological and molecular analysis of three mouse lines that express a tamoxifen-inducible *cre* recombinase to activate *KRAS^G12D^* in acinar cells of the pancreas. The cre-recombinase was targeted to the acinar-specific transcription factor genes, *Ptf1a* and *Mist1/Bhlha15*, or expressed within a BAC-derived *Elastase* transgene. Up to two months after tamoxifen induction of *KRAS^G12D^*, morphological changes were negligible. However, induction of pancreatic injury by cerulein resulted in stark differences in tissue morphology between lines within seven days, which were maintained for at least five weeks after injury. *Ptf1a^creERT^* pancreata showed widespread PanIN lesions and fibrosis, while the *Mist1^creERT^* and *Ela-creERT* models showed reduced amounts of pre-neoplastic lesions. RNA-seq analysis prior to inducing injury suggested *Ptf1a^creERT^* and *Mist1^creERT^* lines have unique profiles of gene expression that predict a differential response to injury. Multiplex analysis of pancreatic tissue confirmed different inflammatory responses between the lines. These findings suggest understanding the mechanisms underlying the differential response to *KRAS^G12D^* will help in further defining the intrinsic KRAS-driven mechanisms of neoplasia initiation.

## Introduction

Pancreatic Ductal Adenocarcinoma (PDAC) is the 3^rd^ leading cause of cancer-related deaths in United States. At ∼12%, PDAC has the poorest five-year survival rate among the major cancers in United States, which has changed minimally in the past four decades (1). Genetic alterations are found in most PDAC tumours, with >90% of patients harboring somatic mutations in *KRAS*. The most common KRAS mutation causes a glycine to aspartic acid change at amino acid position 12 (*KRAS^G12D^*), resulting in constitutive activation of KRAS. Oncogenic *KRAS* mutations are observed in pancreatic intraepithelial neoplasias (PanIN) or acinar to duct cell metaplasia (ADM), which are precursor lesions to PDAC (2). However, in vivo studies in mice support a model in which *KRAS^G12D^*requires additional genetic mutations in tumor suppressors, such as *p53*, *Ink4a*, or *Smad4,* or environmental events, such as pancreatic injury, to progress to PDAC (3).

To understand the cellular and molecular mechanisms that contribute to *KRAS^G12D^*-mediated PDAC progression, genetically engineered mouse models (GEMMs) have been generated that allow spatial and temporal induction of *KRAS^G12D^* (4–9). A wide range of GEMMs exist, recapitulating the key features and mutations seen in human PDAC, including those allowing activation of *KRAS^G12D^* in combination with the inactivation of *p16*, *p53* or *Smad4*, or with hereditary *Brca1* and *Brca2* mutations (4,10). While initial models focused on generating mutations in key genes during embryonic development, more recent GEMMs have focused on inducible, acinar-specific expression of *KRAS^G12D^*, which is more representative of the spontaneous mutations seen in patients (11–14). The most commonly used GEMMs involve targeting a tamoxifen-inducible *cre recombinase* to acinar-specific genes, such as *Ptf1a* (*Ptf1a^creERT/+^LSL-KRAS^G12D^)* and *Mist1/Bhlha15* (*Mist1^creERT/+^LSL-KRAS^G12D^*). *Ptf1a* and *Mist1* (*Bhlha15*) are basic helix-loop-helix (bHLH) transcription factors involved in pancreatic development. PTF1A is required for pancreatic formation and specification of acinar cells during pancreatic organogenesis in developing embryos (15,16). Mice homozygous for a null *Ptf1a* mutation survive embryonic development but die shortly after birth (7). *Mist1* is expressed shortly after *Ptf1a* during embryogenesis and promotes acinar cell maturation (17). *Mist1*-null mice are viable but present extensive disorganization of the exocrine pancreas and experience increased tissue damage over time (18,19). *Mist1* has been described as a scaling factor which, rather than initiating transcription of developmental genes, enhances expression of genes required for acinar cell specialization (17,19,20). In the adult pancreas, *Ptf1a* and *Mist1* help maintain the differentiated acinar cell state by regulating expression of genes producing digestive enzymes or regulating exocytosis (15,21,22). Deletion of either transcription factors in adults has been linked to promotion of pancreatic diseases, including PDAC (16,23,24).

Inducible models of PDAC targeting creERT to *Ptf1a* and *Mist1* are functional heterozygotes. However, loss of a single *Ptf1a* gene affects complete acinar cell differentiation. Ptf1a heterozygotes have enhanced PDAC progression when KRAS^G12D^ is expressed in the developing pancreas (16,25). Whether this difference in sensitivity occurs in the inducible models and affects PDAC progression in adult acini remains an open question. The goal of this study was to compare the effects of KRAS^G12D^ activation in the inducible GEMMs. Morphological and molecular responses to KRAS^G12D^ activation were compared with and without pancreatic injury between mice with a single (*Ptf1a^creERT/+^KRAS^G12D/+^*) or two copies (*Mist1^creERT/+^KRAS^G12D/+^* or *Ela-creERT; KRAS^G12D/+^*) of *Ptf1a*. These experiments show a more dramatic response to *KRAS^G12D^* in *Ptf1a* heterozygotes based on extensive PanIN progression, increased immune infiltration, and an overall phenotype that promotes conditions supportive of KRAS^G12D^-dependent transformation. These findings confirm *Ptf1a* heterozygosity increases sensitivity to KRAS^G12D^ activation even when limited to adult acinar cells.

## Materials and Methods

### Mouse Models

All mice experiments were approved by Animal Care Committee at the University of Western Ontario (2020-157; 2020-158) and the University of Michigan (PRO00009814). Mice at the University of Western Ontario were bred into a C57/Bl6 background and harbored an inducible *loxP-stop-loxP-Kras^G12D^*(*KRAS^LSL-G12D^*) within the endogenous *Kras* gene. These mice were crossed to mice expressing an inducible cre recombinase (creERT) targeted to either the *Ptf1a* gene (*Ptf1a^creERT/+^KRAS^G12D/+^*) or the *Mist1/Bhlha15* gene (*Mist1^creERT/+^KRAS^G12D/+^*) to ensure inducible acinar-specific expression of KRAS^G12D^. These lines resulted in a functional heterozygote for PTF1a or MIST1 expression. Mice at the University of Michigan were bred into mixed background mice bearing tdTomato (Ela-CreERT) or C57/Bl6 mice bearing YFP (*Ptf1a^creERT/+^*) in the Rosa 26 locus, with and without the *Kras^LSL-G12D^* allele. The Bac-*Ela-creERT* (*Ela-* creERT) mice contained a BAC-derived transgene with the *Elastase* promoter driving creERT expression (*Ela-creERT; KRAS^G12D/+^*). To induce recombination, 2-4 months old mice were orally gavaged with 3 x 2 (*Mist1^creERT/+^KRAS^G12D/+^*) or 5 x 5 mg (*Ptf1a^creERT/+^KRAS^G12D/+^*) of tamoxifen (TX; Sigma-Aldrich, T5648-5G) at Western University and 5 x 4 mg (*Ela-creERT; KRAS^G12D/+^*and *Ptf1a^creERT/+^KRAS^G12D/+^*) at the University of Michigan over 5 days. This results in >90% recombination for all models (26–28). Mice were either sacrificed 22 days after initial TX gavage or treated with cerulein 1-2 weeks following TX gavage. For cerulein-induced pancreatitis (CIP), mice at Western University received intraperitoneal injections of 50-75 µg/kg cerulein (MedChemExpress, HY-A0190/CS-5876) 8 times over 7 hours, which was repeated after two days. Mice at the University of Michigan received 8 intraperitoneal injections of 75 ug/kg cerulein over 7 hours for two consecutive days. CIP-treated mice were sacrificed 1, 3 or 5 weeks after initial CIP treatment.

### Immunohistochemistry (IHC) and Multiplex Immunostaining (Opal)

At the time of sacrifice, pancreata were dissected and prepared for histological, biochemical, or molecular analysis. Pancreatic tissues were fixed in 4% formalin at 4°C overnight, then embedded in paraffin and sectioned to 5 μm. Formalin-fixed paraffin embedded (FFPE) pancreatic tissue sections were deparaffinized and stained for H&E histology or prepared for IHC or immunofluorescence (IF). For IHC, sections were permeabilized with 0.2% Triton-X in PBS, then blocked for 1 hour in sheep serum with PBS. Following blocking, sections were incubated overnight at 4°C in primary antibodies diluted in blocking solution. Primary antibodies included rabbit α-amylase (1:1000; Abcam, ab21156) and rabbit α-CK19 (1:1000; Abcam, ab52625). Sections were rinsed and incubated for 30 minutes with α-rabbit IgG secondary antibody (1:1000; Vector labs, PK-4001) diluted in blocking solution. For signal detection, the ImmPACT® DAB Substrate Kit (HRP) was used (Vector Laboratories, SK-4105). Images were captured using a Leica Microscope DM5500B DFC365 FX camera with LAS V4.4 software. Positive staining was determined as a percentage of the whole tissue area using ImageJ software.

Alternatively, serial sections (4 µm thick) of FFPE pancreata were stained on the Discovery Ultra XT autostainer (Ventana Medical Systems Inc, Tucson, AZ) for CK19 (1:200) (TROMAIII, University of Iowa hybridoma core) or amylase (1:1500) (A8273, Sigma Aldrich, St Louis, Mo) and counterstained with Mayer’s hematoxylin to differentiate between recovered and transformed areas of the pancreas after CIP. Slides were scanned on the Panoramic Scan Digital Slide Scanner (Perkin Elmer) and analyzed for percentage area of positive stain in the QuPath software v0.4.2 (29).

For multiplex analysis, FFPE tissue slides were rehydrated in triplicate with xylene, followed by a single submersion in 100% ethanol, 95% ethanol, and 70% ethanol. Rehydrated slides were washed in neutral buffered formalin and rinsed with deionized water. Antigen retrieval was performed using sodium citrate (pH 6.0) and tris-EDTA (pH 9.0) buffers. Tissue was blocked using endogenous peroxidase at room temperature, followed by 10% donkey serum overnight at 4°C. Primary antibodies for CD3 (1:100, Abcam, Ab11089), CD8 (1:200, Cell Signaling, 98941), F4/80 (1:500, Cell Signaling, 70076), and CK19 (1:400, DSHB, AB 2133570) were diluted in 1% BSA block and sections incubated for 1 hour at RT. After washing with TBST in triplicate, tissues were incubated with secondary antibody, Opal Anti-Ms + Rb HRP (Akoya Biosciences, Marlborough, MA ARH1001EA), before conjugating with Opal fluorophores Following incubation, tissue samples were rinsed in deionized water and mounted with DAPI ProLong^™^ Diamond Antifade mountant (Thermo Fisher Scientific). Images were taken using the Vectra® Polaris™ work station (Akoya Biosciences).

Multiplex immunofluorescence images were acquired using the Mantra™ Quantitative Pathology work station (Akoya Biosciences). The resulting multispectral slide scan bands (MOTiF) were used for image capture of DAPI, Opal 480 (F4/80), Opal 520 (CD3), Opal 570 (CD8), and Opal 690 (CK19). Slides were scanned and quality of staining assessed after multiplex IHC staining using the Phenochart software (v1.1.0, Akoya Biosciences). The images were then exported for further analysis into QuPath (v0.4.3). For each image, regions of interest of 1 mm2 were tiled over the tissue to cover >90% of the tissue and exclude peripancreatic fat and any peripancreatic or intraparenchymal lymph nodes. Initial cell detection in each image was done using in the DAPI channel (settings – pixel size 0.5mm, background radius 8mm, median filter 0mm, sigma 1.5mm, minimal area 10mm2, maximum area 400mm2, threshold 10, split by shape – YES, cell expansion 3mm, cell nucleus included). Spot check of 10% of ROIs in each images confirmed consistent selection of all cells in all regions. Annotation of identified cells was then performed using either a threshold classifier approach using the cell/cytoplasm maximum marker fluorescence (CD3, CD8, CK19) or the random decision tree machine learning approach within QuPath (F4/80). The classifier leveraged the significantly different morphology of myeloid cells and pancreatic acinar cells and was trained using both fluorescence and morphologic cell characteristics for the classification. Manual spotcheck of all classifiers across multiple ROI in all images revealed <5% of incorrect cell classification using these approaches. The primary issue arose due to compact packing of cells in some areas of the tissue, where fluorescent bleedthrough from a cell affected phenotype classification of its neighbor yielding incompatible phenotypes such as CD3+/F4/80+ cells or CD3+/CK19+ cells. These cases were rare and after manual curation, appropriate phenotypes were able to be assigned in vast majority of these cells. The final phenotype quantification of all cells in each ROI for each image was exported into text data files. Final statistical analysis and data visualization was performed in R (v 4.2.1, packages – tidyverse 1.3.2, pheatmap 1.0.12, RColorBrewer 1.1-3, viridis 0.6.2). Statistical analysis was done with a Kruskal-Wallis rank sum test and subsequent pairwise Wilcoxon rank sum test with p<0.05 considered to be significant.

### RNA isolation, RNA-seq and data analysis

RNA was isolated from pancreatic tissue 22 days after initial TX gavage using Trizol (ThermoFisher Scientific, 15596018) and the PureLink RNA mini kit (ThermoFisher Scientific, 12183018A). RNA quality was confirmed using an Agilent 2100 Bioanalyzer and sequenced to a depth of 50 million reads/sample using the Illumina NextSeq High Output 150 cycle (paired-end sequencing) sequencing kit. For RNA-seq analysis, sequence quality was assessed using FastQC, reads were mapped to the mm10 genome using STAR 2.7.9a (30), counted using the featureCounts option in the Subread aligner v2.0.3 (31), and analyzed for differential expression using DESeq2 package v1.38.3 (32) in RStudio. Batch effect correction and data normalization was performed through built-in DESeq2 features. KEGG and GO pathway analyses were performed using clusterProfiler v4.6.2 R package (33), Gene Set Enrichment Analysis (GSEA) was done using fgsea v1.24.0 R package (34), and heatmaps generated using pheatmap v1.0.12 R package (35). PCAtools 2.10.0 (36) and enrichplot v1.18.4 (37) were used for generating PCA plots and dotplots. Gene sets used for enrichment analysis and visualization were obtained from MSigDB (38,39) and Harmonizome (40). Differentially expression gene (DEG) were defined as any gene with a p-value < 0.05 and fold change (FC) > 2. A threshold of adjusted p-value ≦ 0.05 was set for all pathway analyses.

## Results

Previous studies suggested loss of a single *Ptf1a* allele alters the molecular landscape of acinar cells (16). To determine if this affects the response to oncogenic KRAS (KRAS^G12D^) in adult acinar cells, we compared the morphology of pancreatic tissue 22 days after induction of KRAS^G12D^ in 2 to 4-month-old mice in which creERT was targeted to either the *Ptf1a* or *Mist1* gene (Figure 1A, B), which renders the single floxed allele non-functional. *Ptf1a^creERT/+^KRAS^G12D/+^* and *Mist1^creERT/+^KRAS^G12D/+^* pancreata showed normal pancreatic morphology with no pre-neoplastic lesions. Therefore, we initiated CIP 10 days after the last tamoxifen (TX) treatment to promote pancreatic injury (Figure 1C). Mice received two days of cerulein compared, and compared morphology was compared five weeks after initial cerulein treatment. H&E histology of*_Ptf1a_creERT/+_KRAS_G12D/+* mice revealed an almost complete loss of acinar tissue with extensive regions of low (PanIN2) or high (PanIN3) grade lesions, consistent with our previous work (27). Conversely, significantly fewer PanIN lesions and more acinar tissue was observed in *Mist1^CreERT/+^KRAS^G12D/+^* mice (p < 0.0005, Figure 1D, E). To confirm the differential maintenance of acinar tissue, IHC for markers of acinar (amylase) and duct (CK19) cells was performed. *Ptf1a^creERT/+^KRAS^G12D/+^* pancreata showed significantly more CK19 staining (21.66 ± 3.12%) than *Mist1^creERT/+^KRAS^G12D/+^*tissue (3.94 ± 3.12%, p < 0.005), and minimal amylase staining (0.65 ± 12.89%, p < 0.05) compared to *Mist1^creERT/+^KRAS^G12D/+^*tissue (66.52 ± 12.89%), confirming increased transformation occurred in the *Ptf1a^creERT/+^KRAS^G12D/+^* mice (Figure 1F-H).

**Figure 1.**
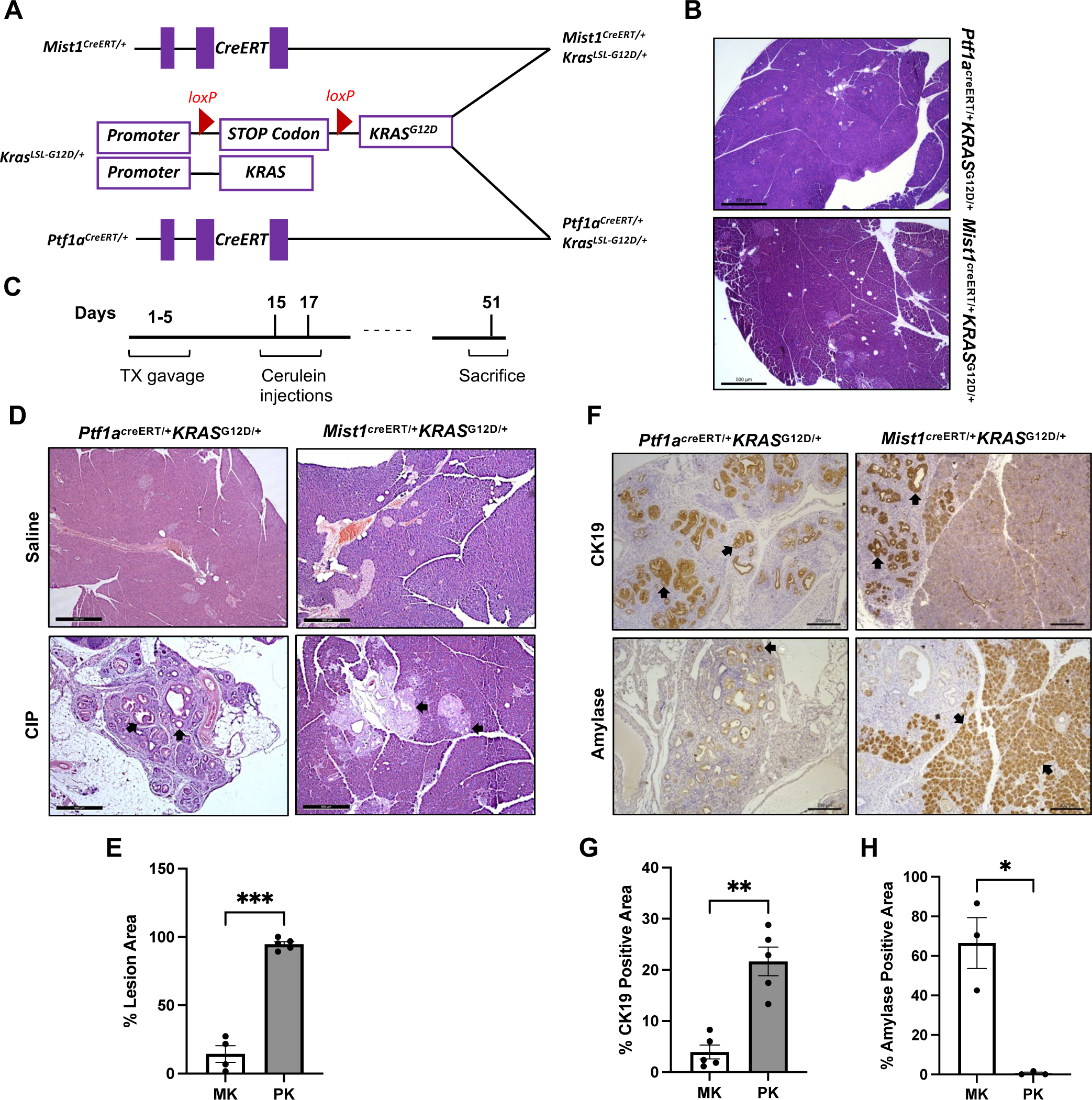
KRAS^G12D^ activation promotes increased damage in a *Ptf1a* haploinsufficient background in response to pancreatic injury. **(A)** Schematic representation of creERT targeting to create inducible knock-in *Ptf1a^creERT/+^KRAS^G12D/+^* (PK) and *Mist1^creERT/+^KRAS^G12D/+^* (MK) mouse lines. **(B)** Representative H&E histology shows no difference between pancreatic morphology of *KRAS^G12D/+^* mice in *Ptf1a^creERT/+^* or *Mist1^CreERT/+^* cohorts 22 days following tamoxifen treatment. Magnification bar=100 µm. **(C)** Experimental timeline for tamoxifen and cerulein treatments. **(D)** H&E histology and **(E)** quantification of percent lesion area show more lesions formed in *Ptf1a^creERT/+^KRAS^G12D^* compared to *Mist1^creERT/+^Kras^G12D^* mice 5 weeks after cerulein treatment. Magnification bar= 500 µm. **(F)** IHC for amylase or CK19, and quantification of the percent **(G)** CK19 or **(H)** amylase positive area in *Ptf1a^creERT/+^KRAS^G12D^* and *Mist1^creERT/+^KRAS^G12D^* mice 5 weeks after cerulein treatment. Magnification bar= 200 µm. In all cases, *\*p* < 0.05, *\*\*p* < 0.005, *\*\*\*p* < 0.0005. Significance was determined using a two-tailed unpaired t-test with Welch’s correction. Error bars represent mean ± standard error. Each dot represents an individual mouse.

To determine if the loss of a single *Ptf1a* allele was the cause of a different response to KRAS^G12D^, *Ptf1a^creERT/+^KRAS^G12D/+^*mice bred in a different institute were compared to a second mouse model that retains two copies of the *Ptf1a* gene. In these mice, KRAS^G12D^ is induced in a transgenic mouse model in which creERT is expressed under the control of the *Elastase* promoter within a bacterial artificial chromosome construct (BAC). In the absence of cerulein, neither mouse line showed pre-neoplastic lesions, similar to earlier findings (Figure 2A). Once again, initiation of CIP in these mice 7 days after induction of *KRAS^G12D^* resulted in widespread PanINs in *Ptf1a^creERT/+^KRAS^G12D/+^* mice extending through the entire tissue (Figure 2B, C). Similar to the *Mist1^creERT/+^KRAS^G12D/+^*mice, *Ela-creERT; KRAS^G12D/+^* tissue showed less CK19 IHC staining indicative of fewer PanINs (1.42 ± 0.606%) relative to *Ptf1a^creERT/+^KRAS^G12D/+^* tissue (15.9 ± 1.22, p < 0.0001) and retained more acinar cell tissue (62.0 ± 4.97%) compared to the *Ptf1a^creERT/+^KRAS^G12D/+^*tissue (33.5 ± 5.38%, p < 0.0041) as indicated by amylase IHC staining (Figure 2D-F). These findings support a model in which absence of a single *Ptf1a* allele increases the sensitivity to KRAS^G12D^-mediated PanIN progression.

**Figure 2.**
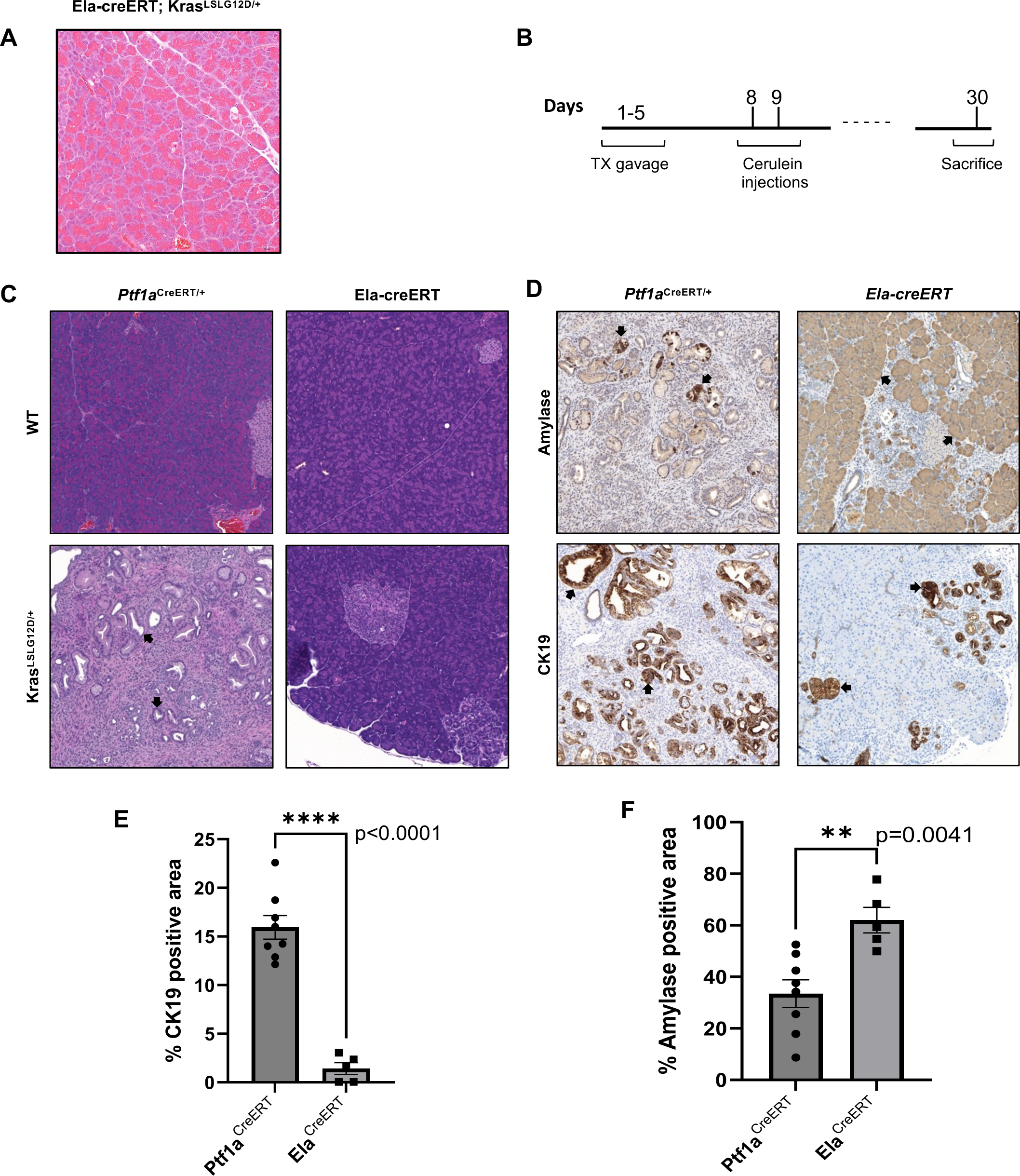
*Ptf1a^creERT/+^* mice show more damage than *Ela-creERT/KRAS^G12D^* in response to KRAS^G12D^ activation following pancreatic injury. **(A)** Representative H&E histology of *Ela-creERT/KRAS^G12D^* mice shows no lesions 22 days following tamoxifen treatment. **(B)** Timeline for tamoxifen and cerulein treatments of the *Ptf1a^creERT/+^KRAS^G12D/+^* and *Ela-creERT/KRAS^G12D^* mice. **(C)** Representative H&E histology shows more lesions in *Ptf1a^creERT/+^Kras^G12D^* mice compared to *Ela-creERT/KRAS^G12D^* mice 3 weeks after cerulein treatment. **(D)** IHC for amylase or CK19 and quantification of the % of tissue area positive for **(E)** CK19 or **(F)** amylase show increased CK19 and decreased amylase staining in *Ptf1a^creERT/+^KRAS^G12D^* mice compared to *Ela-creERT/KRAS^G12D^* pancreata 3 weeks after cerulein treatment. *\*\*p* < 0.01, *\*\*\*\*p* < 0.0001. Significance was determined using a two-tailed unpaired t-test. Error bars represent mean ± standard error. Each dot represents an individual mouse. Magnification bars= 50 uM

To determine if the increased transformation observed in *Ptf1a* heterozygous mice was due to a differential response to injury (i.e. the acinar cells had differential sensitivity) or a differential ability to recover (i.e. the resulting ADM lesions had differential sensitivity), pancreatic morphology was examined 7 days after the initial CIP treatment in all lines (Figure 3). Consistent with longer time points, *Ptf1a^creERT/+^KRAS^G12D/+^*mice showed extensive loss of acinar tissue with widespread PanIN lesion formation (Figure 3B, D), while *Mist1^creERT/+^KRAS^G12D/+^* (Figure 3B) and *Ela^creERT^KRAS^G12D/+^*mice (Figure 3D) retained higher amounts of acinar tissue suggesting loss of a single *Ptf1a* allele increases sensitivity to KRAS^G12D^ and is not simply preventing regeneration.

**Figure 3.**
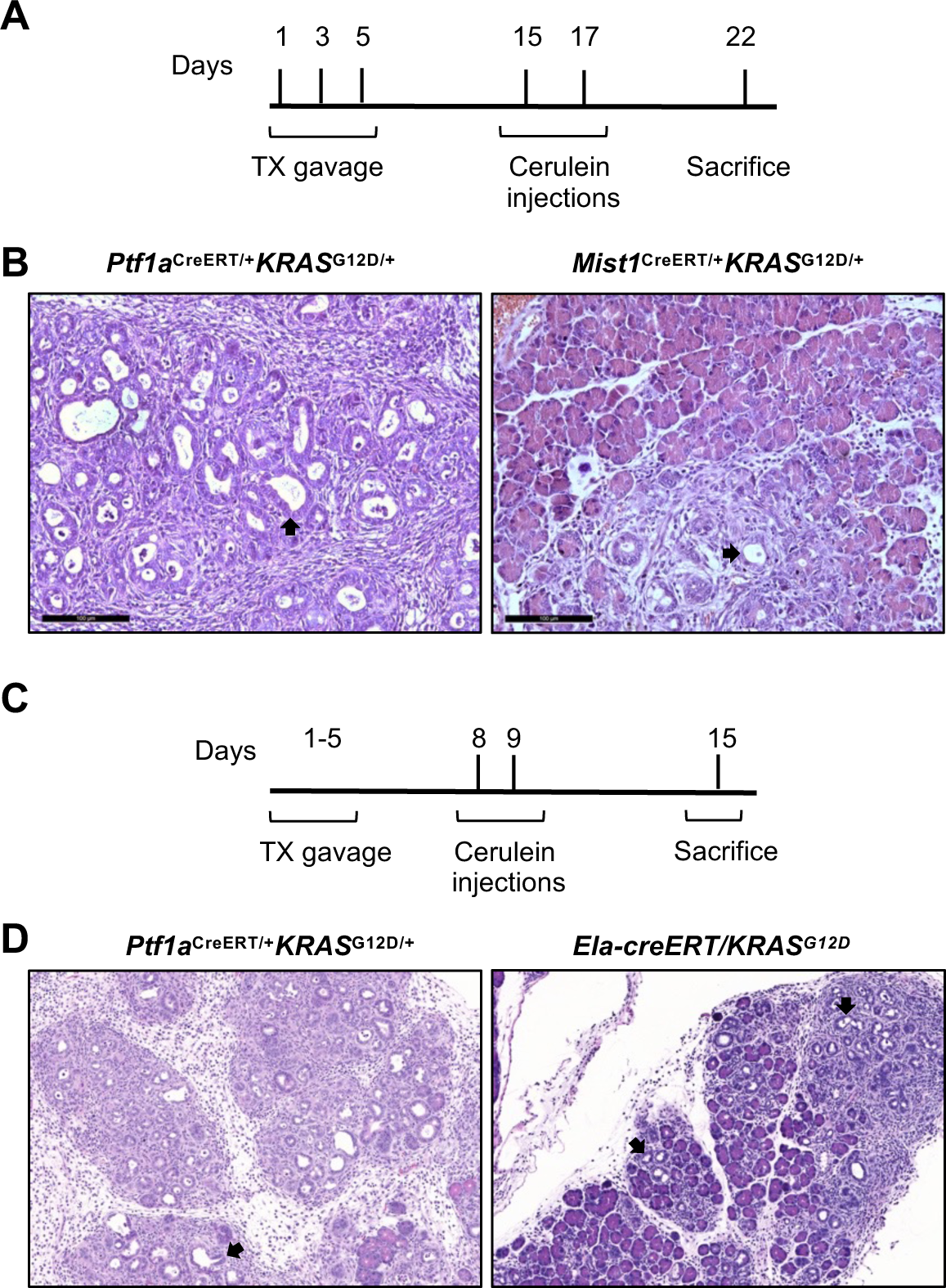
The absence of one *Ptf1a* allele increases loss of the acinar cells in response to KRAS^G12D^ and injury. Timeline of tamoxifen (TX) and cerulein treatment for *Ptf1a^creERT/+^KRAS^G12D^*and **(A)** *Mist1^creERT/+^KRAS^G12D^* or **(C)** *Ela-creERT/KRAS^G12D^* mice. Representative H&E histology shows more damage in the pancreas of *Ptf1a^creERT/+^KRAS^G12D^* than **(B)** *Mist1^creERT/+^KRAS^G12D^* or **(D)** *Ela-creERT/KRAS^G12D^* mice 7 days after cerulein treatment. Magnification bar= 500 µm.

Next, to determine the mechanisms underlying this differential response to KRAS^G12D^ between *Ptf1^+/-^* and *Ptf1^+/+^*tissue, RNA-seq was performed 22 days after initial TX treatment on pancreatic tissue obtained from *Ptf1a^creERT/+^KRAS^G12D/+^*and *Mist1^creERT/+^KRAS^G12D/+^* mice in the absence of pancreatic injury (Figure 4). This experimental timeline was chosen since no morphological differences were observed between *Ptf1a^creERT/+^KRAS^G12D/+^*and *Mist1^creERT/+^KRAS^G12D/+^* pancreatic tissue (Figure 1B). Principal Component Analysis (PCA) comparing *KRAS^G12D^*-expressing and WT pancreata of each line showed distinct clustering of the groups based on their genotype (4A, B), suggesting underlying differences at the transcriptomic level even in the absence of any observable histologic difference. Surprisingly, *Mist1^creERT/+^KRAS^G12D/+^* mice have more differentially expressed genes (DEGs) compared to wild type mice (Figure 4C; 784 DEGs, 439 up and 345 down) than *Ptf1a^creERT/+^KRAS^G12D/+^*mice compared to wild type mice (Figure 4D; 461 DEGs, 374 up and 87 down). A higher percentage of the DEGs in the *Ptf1a^creERT/+^KRAS^G12D/+^*-wild type dataset was upregulated indicating involvement of active regulatory processes.

**Figure 4.**
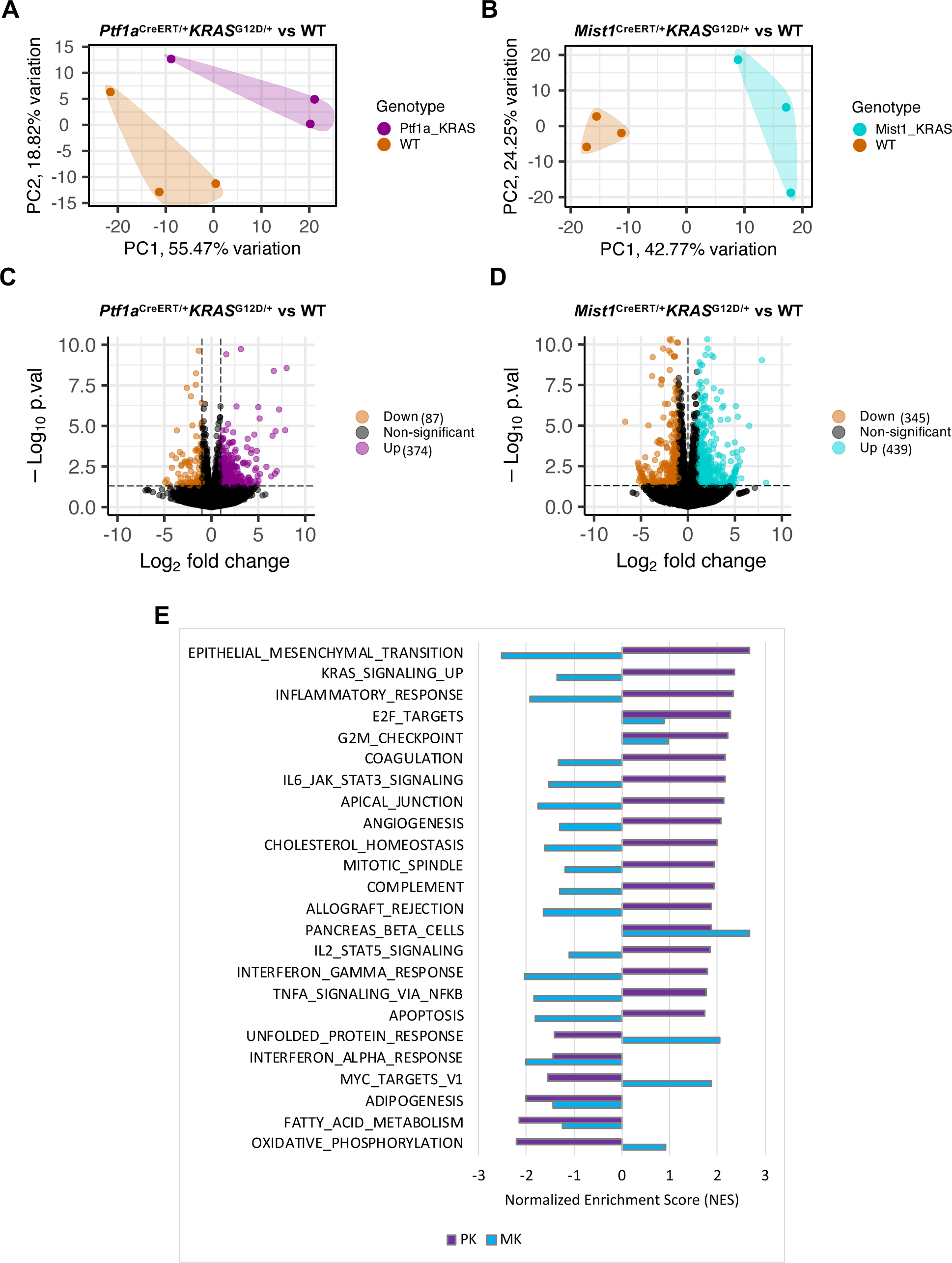
RNA-seq analysis reveals distinct gene expression profiles between *Ptf1a^CreERT/+^Kras^G12D^* and *Mist1^CreERT/+^Kras^G12D^* mice. **(A, B)** PCA plots showing clustering of *KRAS^G12D^* and wild type (WT) RNA from pancreatic tissues of (A) *Ptf1a^creERT/+^KRAS^G12D^* (PK) and (B) *Mist1^creERT/+^KRAS^G12D^* (MK) mice. **(C, D)** Volcano plots comparing differentially-expressed genes (DEGs) between WT and (C) *Ptf1a^creERT/+^KRAS^G12D^* or (D) *Mist1^creERT/+^KRAS^G12D^* RNA-seq datasets. Number of differentially expressed genes are indicated on the graphs. In all cases, DEGs have p-value < 0.05 and log_2_FC> 1. **(E)** Gene Set Enrichment Analysis (GSEA) plot showing normalized enrichment score of the top Hallmark gene sets in *Ptf1a^creERT/+^KRAS^G12D^* and *Mist1^creERT/+^KRAS^G12D^* RNA compared to their corresponding wild type RNA samples.

Next, Gene Set Enrichment Analysis (GSEA) was performed using MSigDB’s hallmark gene sets comparing *Ptf1a^creERT/+^KRAS^G12D/+^*and *Mist1^creERT/+^KRAS^G12D/+^* to their respective wild type control tissues. Multiple cancer-related gene sets were differentially enriched between lines (Figure 4E), including positive enrichment of genes defining epithelial-mesenchymal transition, Kras signaling, and inflammatory response. These differences appeared in the absence of morphological changes in pancreatic tissue suggesting unique molecular gene expression profiles exist between the lines prior to phenotypic transformation.

One of the gene sets identified by GSEA was genes up-regulated by KRAS activation (HALLMARK_KRAS_SIGNALING_UP). This pathway was positively enriched compared to wild type tissue in the presence of one *Ptf1a* allele (*Ptf1a^creERT/+^KRAS^G12D/+^*) but negatively enriched when two *Ptf1a* alleles were present (*Mist1^creERT/+^KRAS^G12D/+^*; Figure 5A). Comparison of normalized *KRAS* read counts from the sequencing files indicates an increase in expression, but these increases were not significant in either the *Ptf1a^creERT/+^KRAS^G12D/+^* - wild type or *Mist1^creERT/+^KRAS^G12D/+^* - wild type comparisons (Figure 5B), and *Kras* was not identified as a DEG in either dataset. This suggests the positive enrichment for KRAS signaling in the *Ptf1a^creERT/+^KRAS^G12D/+^*pancreas is due to upregulation of downstream KRAS effectors rather than differences in *KRAS* expression. Plotting normalized read counts of genes in the MAPK signaling pathway from the KEGG pathways dataset for wild type and *Ptf1a^creERT/+^KRAS^G12D/+^* RNA shows an overall pattern of increased expression for the MAPK pathway in *Ptf1a^creERT/+^KRAS^G12D/+^*mice (Figure 5C). In confirmation of this, IHC for FOS, a member of the Activator Protein 1 (AP-1) transcriptional complex, downstream mediator of KRAS signaling in the MAPK pathway, and identified as a DEG only in *Ptf1a^creERT/+^KRAS^G12D/+^* mice, showed increased accumulation in the *Ptf1a^creERT/+^KRAS^G12D/+^*pancreas compared to *Mist1^creERT/+^KRAS^G12D/+^* tissue, but only after induction pancreatic injury (Figure 5D). JUN, a binding partner for FOS did not show the same increased expression in IHC. This is in line with the RNA-seq data as *Jun*, unlike *Fos*, does not show differential expression of Jun at the transcriptomic level (Figure 5E). Combined, these findings suggest the loss of a single *Ptf1a* allele causes differential enrichment of KRAS signaling, which may account for the more aggressive phenotypes in pancreatic cancer mouse models.

**Figure 5.**
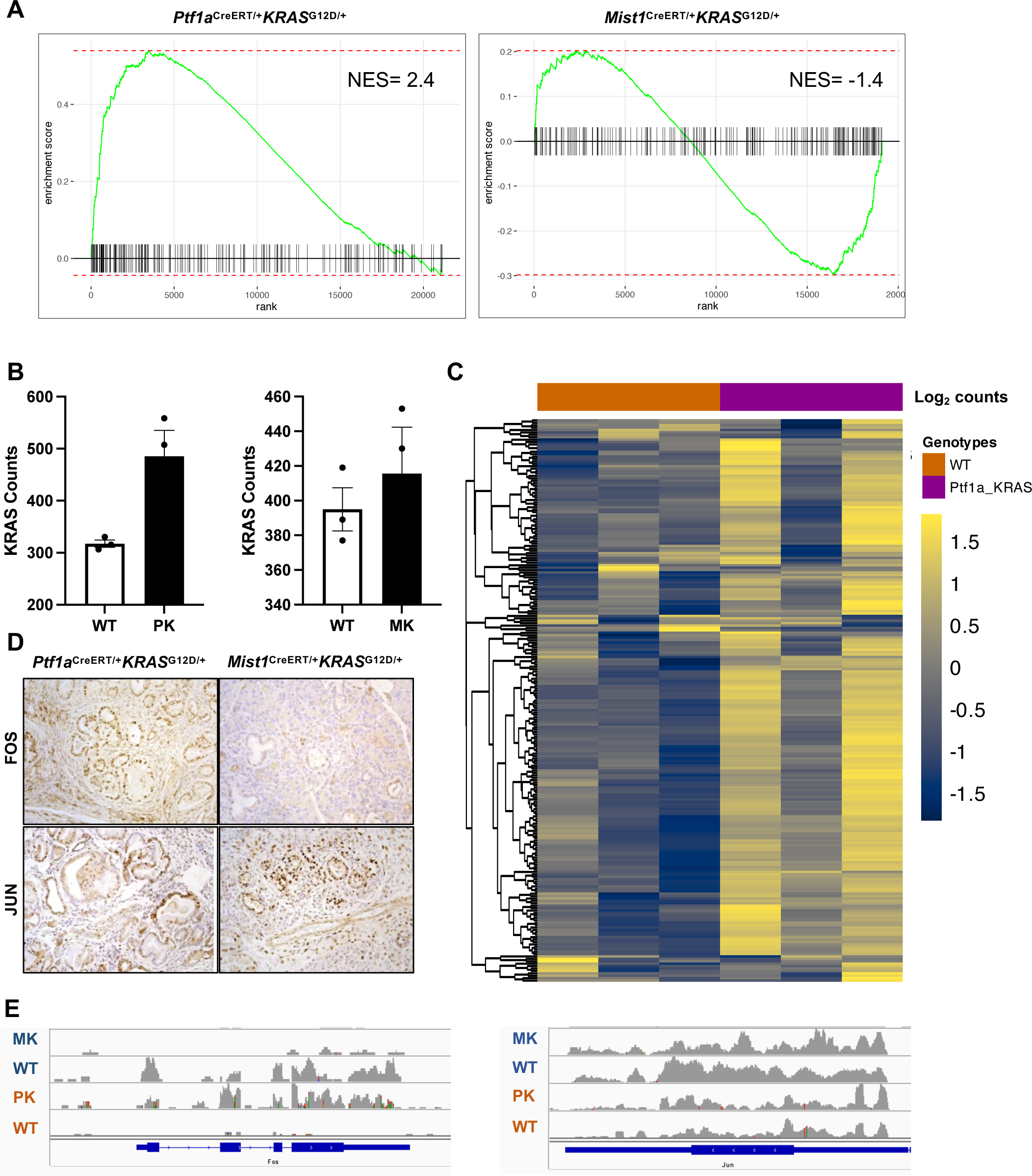
KRAS signalling is enhanced by loss of a single *Ptf1a* allele. **(A)** GSEA plots for “Hallmark genes upregulated by KRAS activation” shows positive enrichment in *Ptf1a^creERT/+^KRAS^G12D^* but not *Mist1^creERT/+^KRAS^G12D^* RNA-seq datasets compared to their corresponding WT mice. **(B)** Bar plot for normalized read counts of *KRAS* expression based on *Ptf1a^creERT/+^KRAS^G12D^* (PK) and *Mist1^creERT/+^KRAS^G12D^* (MK) RNA-seq datasets show no difference in *KRAS* expression compared to wild type (WT). Significance was determined using a two-tailed unpaired t-test with Welch’s correction. Error bars represent mean ± standard error. N values are indicated on top of each bar. **(C)** Heatmap for Log_2_ of normalized read counts of genes in the KEGG pathway for MAPK Signaling shows increased expression of these genes in the *Ptf1a^creERT/+^KRAS^G12D^* mice compared to WT. **(D)** IHC for FOS (upper panels) and JUN (lower panels) shows more positive nuclei in *Ptf1a^CreERT/+^Kras^G12D^* tissue compared to *Mist1^creERT/+^KRAS^G12D^* tissue for FPS but not JUN. Magnification bar= 500 µm. **(E)** RNA tracks showing expression level of Fos (left) and Jun (right) in the *Mist1^creERT/+^KRAS^G12D^* tissue and its corresponding wild type and the *Ptf1a^creERT/+^KRAS^G12D^* tissue and its corresponding wild type tissue.

Next, the molecular profile of the two lines were assessed by Gene Ontology (GO) pathway analysis. 154 GO pathways were enriched based on the *Mist1^creERT/+^KRAS^G12D/+^* - wild type comparison set, while 403 pathways were enriched based on the *Ptf1a^creERT/+^KRAS^G12D/+^* - wild type comparison set. However, only 15 pathways were common between these two comparisons. The top pathways uniquely enriched in the *Ptf1a^creERT/+^KRAS^G12D/+^*-wild type comparison set were almost exclusively immune-related (Figure 6A, B), similar to the GSEA findings which indicated hallmark genes in the Inflammatory Response gene set are differentially enriched between *Ptf1a^creERT/+^KRAS^G12D/+^*and *Mist1^creERT/+^KRAS^G12D/+^* tissues (Figure 4E). GSEA shows significantly positive enrichment in the *Ptf1a^creERT/+^KRAS^G12D/+^*-wild type comparison set (NES= 2.4; Figure 6c) and significant negative enrichment *Mist1^creERT/+^KRAS^G12D/+^* - wild type comparison set (NES= -1.9). These results predict a differential inflammatory response between *Ptf1a^creERT/+^KRAS^G12D/+^*and *Mist1^creERT+^KRAS^G12D/+^* mice.

**Figure 6.**
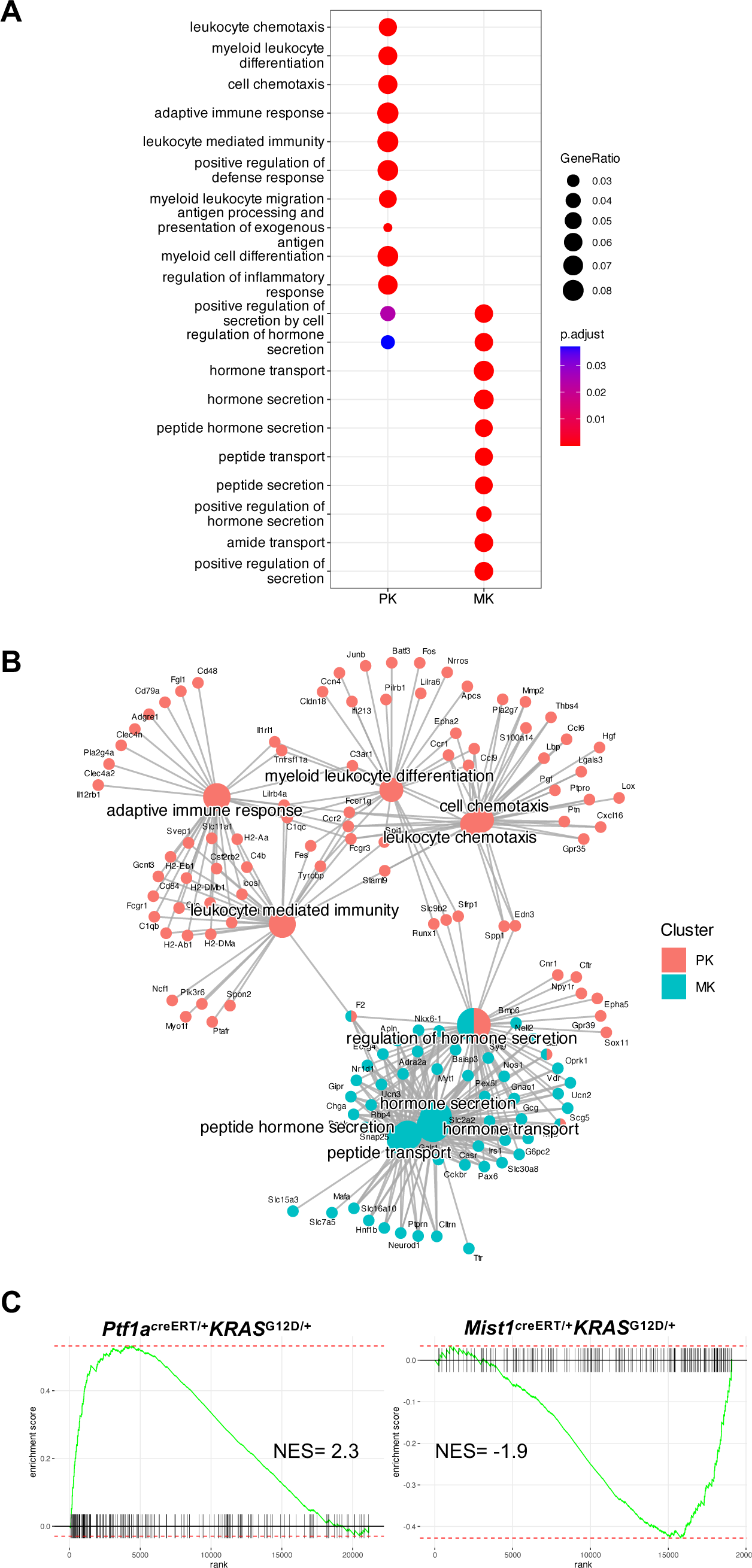
A distinct inflammatory response is activated in the *Ptf1a^CreERT/+^Kras^G12D^*model. **(A)** Dotplot and **(B)** Cnet plot shows top Gene Ontology (GO) pathways enriched in *Ptf1a^creERT/+^KRAS^G12D^* (PK) and *Mist1^creERT/+^KRAS^G12D^* (MK) tissue. Immune-related terms are over-represented in the *Ptf1a^creERT/+^KRAS^G12D^* dataset but not in *Mist1^creERT/+^KRAS^G12D^*. The analysis was performed using DEGs with a p-value <0.05 and log_2_FC> 1. **(C)** GSEA plots for Hallmark “Genes in the Inflammatory Response” in the MSigDB database shows positive enrichment scores for *Ptf1a^creERT/+^KRAS^G12D^*but not *Mist1^creERT/+^KRAS^G12D^* compared to the corresponding wild type cohorts.

To examine immune cell infiltration following CIP in the two mouse lines, we performed an Opal Multiplex assay for multiple immune marks in *Ptf1a^creERT/+^KRAS^G12D/+^*and *Mist1^creERT+^KRAS^G12D/+^* mice one week after CIP (Figure 7). CK19 IF showed reduced staining in *Mist1^creERT+^KRAS^G12D/+^*tissue compared to *Ptf1a^creERT/+^KRAS^G12D/+^* (Figure 7A, B) consistent with earlier analysis. F4/80 accumulation marks macrophages and is generally correlated with the level of CK19 staining observed in the tissue. *Ptf1a^creERT/+^KRAS^G12D/+^*pancreata showed the highest load of macrophage infiltration around lesions (average percent per tile= 25.8%; Figure 7A, C). CD8+ T cells (average percent per tile= 0.18%; Figure 7A, D) were significantly lower in number compared to macrophages, but still showed significantly more accumulation in *Ptf1a^creERT/+^KRAS^G12D/+^*tissue compared to *Mist1^creERT+^KRAS^G12D/+^* tissue. CD3^+^ T cells (average percent per tile= 0.04%; Figure 7A, E) did not show any significant difference between the two lines. Hierarchical clustering of the expression level of each immune marker in *Ptf1a^creERT/+^KRAS^G12D/+^*, *Mist1^creERT+^KRAS^G12D/+^*, and wild type tissues following quantification confirmed a higher percentage of lesions (CK19) and macrophage infiltration (F4/80) in the *Ptf1a^creERT/+^KRAS^G12D/+^*tissue (Figure 7F) suggesting a correlation between CK19 and F4/80 staining (Figure 7G) consistent with the observed macrophage infiltration around CK19+ ADM and PanINs.

**Figure 7.**
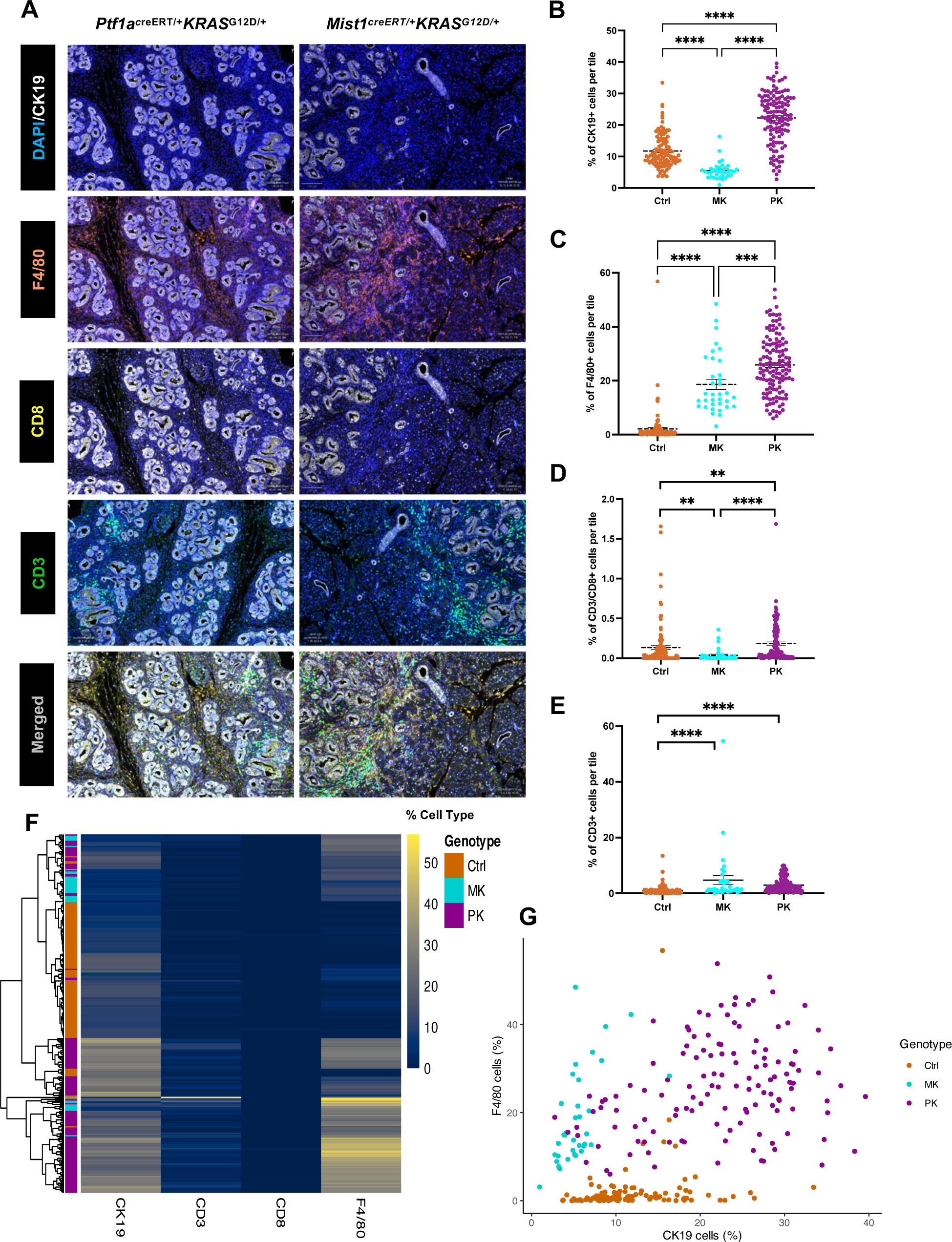
Macrophage and CD3+ T-cell infiltration is increased in the *Ptf1a^creERT/+^KRAS^G12D^* pancreas. **(A)** Opal Multiplex assays for CK19 and multiple immune marks, including F4/80 (macrophage marker), CD8+ (cytotoxic) T-cells, and CD3^+^ T-cells show increased CK19, F4/80 and CD3+ accumulations in the *Ptf1a^creERT/+^KRAS^G12D^* (PK) pancreas compared to *Mist1^creERT/+^KRAS^G12D^* (MK) tissue. **(B-E)** Scatter plots displaying the percent of CK19+, F4/80+, CD8+, and CD3+ cells per tile in *Ptf1a^creERT/+^KRAS^G12D^*, *Mist1^creERT/+^KRAS^G12D^*, and wild type (WT) pancreata. In all cases, *\*\*p* < 0.005, *\*\*\*p* < 0.0005, and *\*\*\*\*p* < 0.0001. Significance was determined using Kruskal-Wallis rank sum test and subsequent pairwise Wilcoxon rank sum test. Error bars represent mean ± standard error. **(F)** Heatmap showing expression level and clustering of each immune marker in the wild type (Ctrl), *Ptf1a^CreERT/+^Kras^G12D^* and *Mist1^CreERT/+^Kras^G12D^* tissue, following quantification on the QuPath software. **G)** Scatter plot visualizing percentage of cells stained with F4/80 against those stained with CK19 in WT, *Ptf1a^creERT/+^KRAS^G12D^*, and *Mist1^creERT/+^KRAS^G12D^* tissue.

To determine if the differential immune cell infiltration also occurs between the *Ptf1a^creERT/+^KRAS^G12D/+^* and *Ela-creERT; KRAS^G12D/+^* mouse lines, similar analysis was performed 1 week following CIP (Figure 8). Surprisingly, *Ptf1a^creERT/+^KRAS^G12D/+^* tissue did not show increased CK19+ lesions compared to *Ela-creERT; KRAS^G12D/+^*tissue (Figure 8A, B) even though the number of progressed lesions are more in *Ptf1a^creERT/+^KRAS^G12D/+^*. However, Opal Multiplex assays for F4/80 showed *Ptf1a^creERT/+^KRAS^G12D/+^*pancreata had increased macrophage infiltration (Figure A, C) compared to *Ela-creERT; KRAS^G12D/+^* tissue with no difference observed between the two lines for CD8^+^ and CD3^+^ T cell infiltration (Figure 8A, D, E). Hierarchical clustering of the expression level of each immune marker in *Ptf1a^creERT/+^KRAS^G12D/+^*, *Ela-creERT; KRAS^G12D/+^*, and the wild type tissues following quantification confirmed a higher percentage of F4/80 staining (Figure 8F). The higher level of CK19 staining in the *Ela-creERT; KRAS^G12D/+^* tissue does not seem to be correlated with higher level of macrophage infiltration (Figure 8G), which could suggest that the higher percentage of CK19 positive cells in these mice are a byproduct of normal ducts and recovering stressed acinar cells with residual CK19 signal. There is however a positive correlation between CK19 staining and macrophage infiltration in the *Ptf1a^creERT/+^KRAS^G12D/+^*tissue (Figure 8G). These findings support a model in which immune-related pathways are differentially activated in *Ptf1a^creERT/+^KRAS^G12D/+^* mice. These observations confirm RNA-seq analysis showing differential enrichment of immune-related pathways between lines and that loss of a single *Ptf1a* alleles may prime the tissue for a differential immune response even before appearance of any lesions.

**Figure 8.**
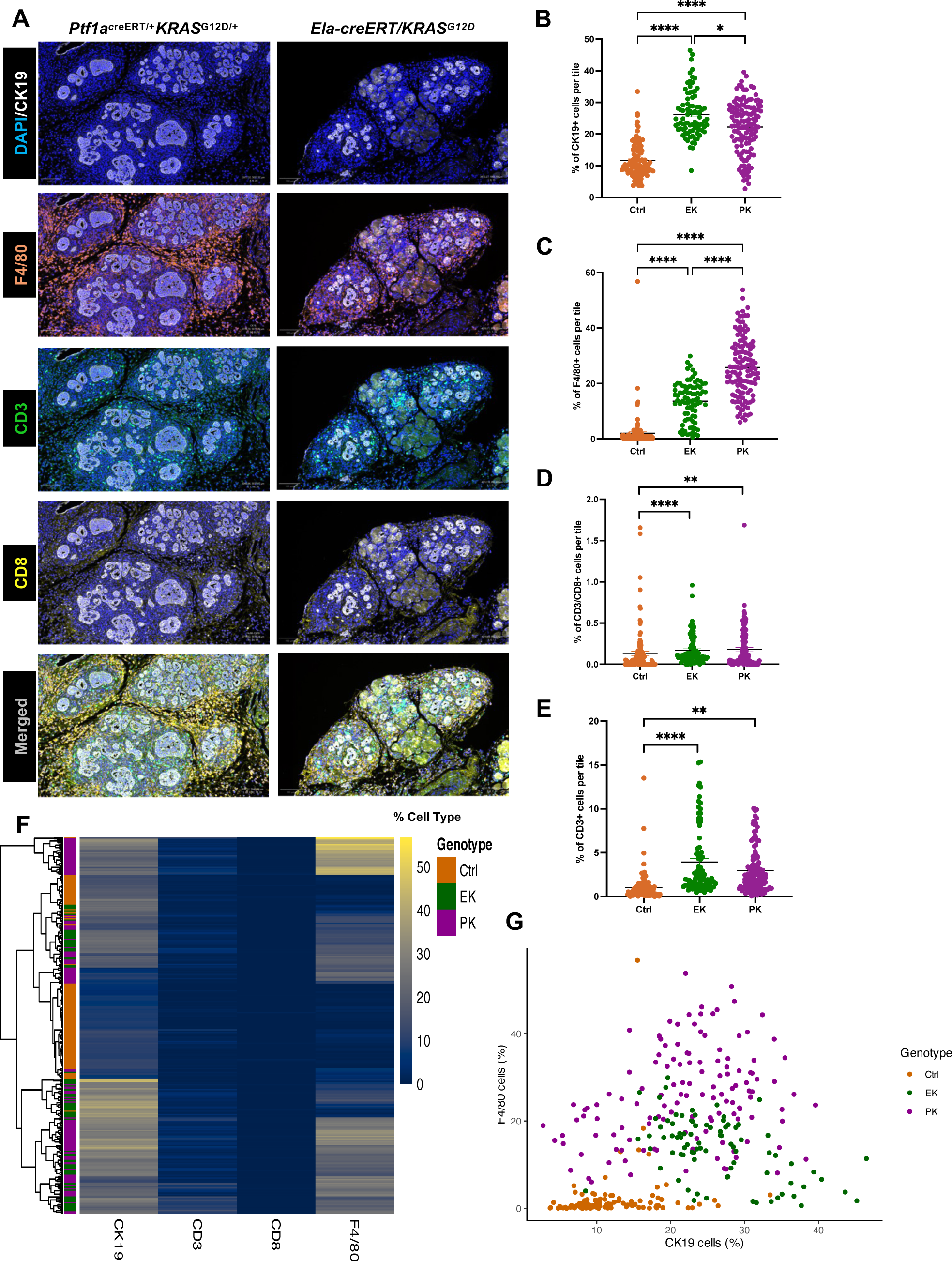
Macrophage and CD3+ T-cell infiltration is increased in the *Ptf1a^creERT/+^KRAS^G12D^* pancreas. **(A)** Opal Multiplex assays for CK19 and multiple immune marks, including F4/80 (macrophage marker), CD8+ (cytotoxic) T-cells, and CD3^+^ T-cells show increased CK19, F4/80 and CD3+ accumulations in the *Ptf1a^creERT/+^KRAS^G12D^* (PK) pancreas compared to *Ela-creERT/KRAS^G12D^* (EK) tissue. **(B-E)** Scatter plots displaying the percent of CK19+, F4/80+, CD8+, and CD3+ cells per tile in *Ptf1a^creERT/+^KRAS^G12D^*, *Ela-creERT/KRAS^G12D^*, and wild type (WT) pancreata. In all cases, *\*p* < 0.05, *\*\*p* < 0.005, *\*\*\*p* < 0.0005, and *\*\*\*\*p* < 0.0001. Significance was determined using Kruskal-Wallis rank sum test and subsequent pairwise Wilcoxon rank sum test. Error bars represent mean ± standard error. **(F)** Heatmap showing expression level and clustering of each immune marker in the wild type (Ctrl), *Ptf1a^CreERT/+^Kras^G12D^* and *Ela-creERT/KRAS^G12D^* tissue, following quantification on the QuPath software. **G)** Scatter plot visualizing percentage of cells stained with F4/80 against those stained with CK19 in WT, *Ptf1a^creERT/+^KRAS^G12D^*, and *Ela-creERT/KRAS^G12D^* tissue.

## Discussion

Pancreatic ductal adenocarcinoma has the highest mortality rate of the major cancers, with few effective therapeutic options available. Significant advances have been made on the fundamental mechanisms in PDAC initiation and progression through the development of GEMMs. Since the development of the KC mice (41), the first conditional mouse model resulting from breeding *Pdx1-cre* or *Ptf1a-cre* mice to *KRAS^LSL-G12D^* mice, additional genetic mouse models of PDAC have been developed, including inducible and non-inducible models, each informing a unique aspect of PDAC. All these models use an oncogenic KRAS mutation (*KRAS^G12D^* or *KRAS^G12V^*) with the goal of generating temporal and/or spatial activation to mimic the etiology of human PDAC more accurately. Differences have been noted between inducible and non-inducible models of PDAC, and there are indications different cre recombinase drivers may have unique phenotypes. However, to date, a direct comparison between lines has not been published. In this study we characterized the molecular and phenotypic differences between *Ptf1a^creERT/+^KRAS^G12D/+^*, *Mist1^creERT/+^KRAS^G12D/+^*, and *Ela-creERT; KRAS^G12D/+^*mouse lines as these models are among the most used tamoxifen-inducible models allowing Kras expression in adult acinar cells. Our results show clear differences in the molecular response to *KRAS^G12D^* that correlates to marked differences between the lines and highlights the importance for choosing the right model for studying different aspects of pancreatic cancer.

Pancreatic Transcription Factor 1a (PTF1a) is a regulator of pancreatic organogenesis, essential for pancreatic determination, exocrine fate decision, acinar cell differentiation, and activation of other key transcription factors (15,42–44). Null mutations in *Ptf1a* lead to absence of exocrine pancreas and is lethal in mice (7). Our results suggest *Ptf1a* haploinsufficiency resulting from targeted insertion of cre recombinase correlated to increased sensitivity to *KRAS^G12D^* indicated by increased PanIN progression and loss of the mature acinar cell phenotype. These findings are consistent with studies suggesting *Ptf1a*’s function is dosage-dependent, with reduced expression leading to pancreatic hypoplasia, diminished expression of exocrine genes, and ultimately maintenance of progenitor fate (45–47). Furthermore, adult acinar cells in *Ptf1a* heterozygous mice proliferate more than the corresponding wild type mice (48) and *Pdx1^cre^KRAS^G12D^* mice harboring a heterozygous deletion of *Ptf1a* show increased PanINs compared to *Ptf1a* wild type counterparts (16). Our RNA-seq results support a dosage-dependent function of *Ptf1a* and show loss of a single allele is sufficient to alter the molecular response to KRAS^G12D^, even in the absence of morphological changes to the pancreas (Figure 4). Our data suggests loss of one *Ptf1a* allele correlates to increased KRAS signaling, which may account for the increased PDAC progression compared to those lines that retain both copies of *Ptf1a*. This is supported by findings in other studies demonstrating loss of *Ptf1a* gene activates KRAS-dependency and that *Ptf1a* prevents KRAS-driven PDAC progression (16,49).

RNA-seq also suggested an enrichment of inflammatory and immune-related pathways in *Ptf1a^creERT/+^KRAS^G12D^* mice compared to *Mist1^creERT/+^KRAS^G12D^* mice even before the induction of injury. Significantly increased accumulations of macrophages in *Ptf1a^creERT/+^KRAS^G12D^* pancreas, particularly around lesion areas, were observed compared to wild type or *Mist1^creERT/+^KRAS^G12D^*pancreas. This finding reinforces a model in which solid tumors, including PDAC tumor cells, trigger inflammation that recruits immunosuppressive leukocytes, such as macrophages, to the lesion site (50,51). This increased infiltration is a response to KRAS^G12D^ that is programmed early in the initiation stages of the disease, as we see a differential transcriptomic profile before appearance of any lesions that supports myeloid cell differentiation and migration in the *Ptf1a^creERT/+^KRAS^G12D^*mice. This observation is supported by findings from Zhang et al. study indicating that depletion of myeloid cells early during PDAC development prevents PanIN progression (52). The same study showed myeloid cells to be required for sustained MAPK signaling during carcinogenesis, which ultimately promotes immune evasion through a mechanism involving EGFR/MAPK-dependent regulation of PD-L1 expression (52). Similarly in our experiment, along with enrichment of myeloid cell-related pathways and increased F4/80 staining, there is a distinct increase in MAPK pathway-related genes (Figure 5) with the loss of a *Ptf1a* allele (*Ptf1a^creERT/+^KRAS^G12D^*) that was not seen when both *Ptf1a* alleles were functional (*Mist1^creERT/+^KRAS^G12D^*and *Ela-creERT; KRAS^G12D/+^*). This finding proposes a multilateral role for dose-dependency of *Ptf1a* in PDAC progression, understanding which not only contributes to enhancing mouse models studying this progression but may also provide a novel therapeutic strategy for PDAC.

While macrophages represent the most dominant immune cells present in the microenvironment in our study, CD3^+^ and CD8^+^ T-cells showing far less accumulation around PanIN lesions. Macrophage recruitment excludes T-cells from the microenvironment (53), as evident by the scarce number of effector T cells in pre-invasive and invasive PDAC (51). While the overall number of CD8+ and CD3+ T-cells in our study is low in both *Ptf1a^creERT/+^KRAS^G12D^*and *Mist1^creERT/+^KRAS^G12D^* pancreata, the number of CD8+ T cells is significant higher in *Ptf1a^creERT/+^KRAS^G12D^*pancreas compared to *Mist1^creERT/+^KRAS^G12D^*. The heightened aggressivity of the *Ptf1a^CreERT/+^* response to *KRAS^G12D^*may cause a more pronounced immune response in these mice. Whether these T-cells show optimal activation and function and whether they maintain their increased numbers compared to the *Mist1^creERT/+^KRAS^G12D^* mice as the disease progresses should be addressed in future studies.

Overall, our results show that even without pancreatic injury or morphological differences, KRAS^G12D^ promoted unique transcriptomic profiles based on PTF1A dosage, including differential activation of KRAS signaling and enrichment of immune-related pathways. The more extensive lesions and increased immune infiltration suggest *Ptf1a* haploinsufficient mice are primed by KRAS^G12D^, possibly through epigenetic reprogramming, for a differential downstream cascade of events following a second environmental stimulus such as inflammation. Epigenetically primed genes are poised to rapidly respond to external cues, a process utilized during development to regulate cellular differentiation and plasticity (54–58), and observed in a number of diseases including neurodevelopmental disorders, immunodeficiencies, and cancers (59–61). Epigenetic reprogramming of poised genes can increase the sensitivity of pancreatic cells undergoing stress to chronic pancreatitis (62), a risk-factor for PDAC. Future studies are necessary to elucidate whether a link between *Ptf1a* haploinsufficiency and epigenetic mechanisms underlie the differential response to KRAS^G12D^ activation.

In summary, this study highlights the effect of *Ptf1a* haploinsufficiency and dose-dependency on the response to KRAS^G12D^ activation in adult acinar cells. Our findings suggest loss of a single *Ptf1a* allele alters the response to KRAS^G12D^ activation in adult acinar cells, leading to more aggressive phenotypes. One underlying mechanism for this altered response appears to involve differentially priming of genes that include, amongst other things, pathways involved in potentiating KRAS signaling and a pro-inflammatory response at early stages even before the appearance of pre-neoplastic lesions. Follow-up studies investigating such epigenetic mechanisms are necessary to further elucidate the effects of *Ptf1a* haploinsufficiency on PDAC progression and the prognostic significance of lower expression levels of *Ptf1a* in PDAC patients.

## Acknowledgements and Funding

The authors wish to acknowledge the ongoing support of several national research funding agencies for this work including the Canadian Institutes of Health Research (MOP#PJT166029), the Cancer Research Society of Canada and the Rob Lutterman Foundation for Pancreatic Cancer Research. This work would not be possible without specific support from a London Regional Cancer Centre Catalyst Grant, co-supported by Mr. Keith Sammit and an internal bridge grant from the University of Western Ontario. FM is supported by Mitacs Accelerate Fellowship.

## Notes

### Competing Interest Statement

The authors have declared no competing interest.

### Summary of Updates

Title updated to clarify; Abstract updated due to typos.

